# Overcoming analytical reliability issues in clinical proteomics using rank-based network approaches

**DOI:** 10.1101/020867

**Authors:** Wilson Wen Bin Goh, Limsoon Wong

## Abstract

Proteomics is poised to play critical roles in clinical research. However, due to limited coverage and high noise, integration with powerful analysis algorithms is necessary. In particular, network-based algorithms can improve selection of reproducible features in spite of incomplete proteome coverage, technical inconsistency or high inter-sample variability. We define analytical reliability on three benchmarks --- precision/recall rates, feature-selection stability and cross-validation accuracy. Using these, we demonstrate the insufficiencies of commonly used Student’s t-test and Hypergeometric enrichment. Given advances in sample sizes, quantitation accuracy and coverage, we are now able to introduce and evaluate Ranked-Based Network Approaches (RBNAs) for the first time in proteomics. These include SNET (SubNETwork), FSNET (FuzzySNET), PFSNET (PairedFSNET). We also introduce for the first time, PPFSNET(samplePairedPFSNET), which is a paired-sample variant of PFSNET. RBNAs (particularly PFSNET and PPFSNET) excelled on all three benchmarks and can make consistent and reproducible predictions even in the small-sample size scenario (n=4). Given these qualities, RBNAs represent an important advancement in network biology, and is expected to see practical usage, particularly in clinical biomarker and drug target prediction.

## Introduction

Mass spectrometry (MS)-based proteomics has become indispensable for many aspects of biological and clinical research. Although MS-based proteomics has advanced significantly in recent years, data reliability issues still persist. The most common setup is the Data-Dependent Acquisition (DDA) platform where eluting peptides from a separation column are selected for fragmentation semi-randomly. This leads to inconsistent quantitative information amongst identified proteins, and presents a severe analytical challenge.

Recent proteomic advancements have led to the new Data-Independent Acquisition (DIA) paradigm, where fragment precursor selection is independent of stoichiometry, leading to more complete datasets (Venable et al. 2004; Carvalho et al. 2010). One instance of DIA is SWATH, where data is captured by repeatedly cycling through precursor isolation windows (SWATH windows) within a predefined m/z range (Gillet et al. 2012).

Guo et al. have shown that, when coupled with PCT (Pressure Cycling Technology), SWATH could be used to reproducibly digitize the proteome of minute amounts of clinical samples in a high-throughput fashion (Guo et al. 2015). Although they demonstrated this approach was feasible, what was noteworthy, and of concern, was also the fact that SWATH is ostensibly noisier due to the concurrent fractionation of a large number of precursors.

A second concern, which is of clinical and biological relevance, is the proteome coverage issue. Although SWATH captures a large amount of spectral data, two major bottlenecks exist: 1/ identification of proteins is still reliant on an incomplete reference spectral library, and 2/ sparse alignments of peak groups against a noisy background (Rosenberger et al. 2014). On average, SWATH can identify between 2,000 to 5,000 proteins (Rosenberger et al. 2014). While this is clearly higher than SRM/MRM assays, a large segment of the proteome and the underlying changes relevant to the phenotype remain unexamined.

Networks can be combined with proteomics synergistically (Gstaiger and Aebersold 2009; Bensimon et al. 2012; Goh et al. 2012a; Goh and Wong 2013; Goh et al. 2013c; Goh and Wong 2014) to overcome its idiosyncratic coverage and consistency issues. At its most primitive, simple cluster prediction based on overlaying DDA-identified differential proteins against a protein interaction network yielded a list of undetected but relevant proteins that can be iteratively re-examined in the raw spectra (Li et al. 2009; Goh et al. 2011; Goh et al. 2012b). A more sophisticated approach uses a list of subnets as a feature vector that can be scored by its overlaps with the set of identified proteins, thus overcoming consistency issues where protein numbers differ sample-to-sample (Goh et al. 2012c; Goh et al. 2013a).

Thus, improving analytical outcome of SWATH via network integration is a tantalizing prospect. Additionally, two aspects of SWATH, as an instance embodying the qualities of next-generation proteomics, afford new opportunities to test more powerful approaches. 1/ SWATH provides absolute quantification values that approximate SRM/MRM accuracy, the current gold standard for proteomic quantification; and 2/ SWATH can be deployed on complex clinical data with reasonable coverage, sample size and consistency. In this work, we examine the applicability and performance of Rank-Based Network Approaches (RBNAs) on proteomics.

Examples of RBNAs include SNET (SubNETworks) and its successors, FSNET (Fuzzy SNET) and PFSNET (Paired FSNET) (Soh et al. 2011; Lim and Wong 2014), which were proposed originally in the context of gene expression profile analysis. Broadly, they work in the following steps. First, features are ranked in inverse order (highest to lowest abundance) for each tissue. A cut-off at a predefined alpha level is used to identify the set of top alpha features for each tissue. Relevant features are the set of alpha features found in at least beta% of the tissues of a class. In the second step, relevant features are used to fragment known pathways into subnets. Thirdly, each subnet is scored on each tissue according to the expression levels of the features constituting the subnet. Class-specific weights are introduced to modulate the scores. And finally, differential subnets are determined in the statistical feature-selection step. We adapt these methods for proteomics profile analysis here. For details, refer to **Material and methods**.

Rank-based methods are powerful: Rank-based approaches have been shown to be more robust than those utilizing full expressional information. Recent evaluations by Patil *et al* (Patil et al. 2015) and Lim *et al* (Lim et al. 2015) demonstrated that the features selected by rank-based approaches are stable and generalizable onto other similar datasets. Building upon this, RBNAs, which utilize rank information, have been shown to be extremely powerful for identification of relevant features, producing unparallelled prediction reliability and reproducibility in transcriptomics studies (Soh et al. 2011; Lim and Wong 2014). However, they have never been studied on proteomics data due to lack of high-quality proteomics datasets with sufficient sample size or reliable absolute quantification.

In this paper, using a recently published SWATH dataset, we benchmark the standard two-sample t-test feature-selection method (Single Protein, or SP), a standard expression-based network analysis approach using the hypergeometric-enrichment (HE) method, three known RBNAs (SNET, FSNET and PFSNET), and a new variant of PFSNET, which incorporates a novel inter-class pairing feature-selection method, called Paired-sample PFSNET, or simply PPFSNET. The reliability benchmarks are based on the summed performance rankings on 3 criteria, viz. 1/ precision-recall rates, 2/ feature-selection stability, and 3/, cross-validation prediction accuracy.

## Material and methods

### SWATH data

The SWATH dataset from (Guo et al. 2015) was used in this study. This dataset contains 24 SWATH runs from 6 pairs of non-tumorous and tumorous clear-cell renal carcinoma (ccRCC) tissues, which have been swathed in duplicates (12 normal, 12 cancer).

## SWATH data interpretation

All SWATH maps were analyzed using OpenSWATH (Rost et al. 2014) and a spectral library containing 49,959 reference spectra for 41,542 proteotypic peptides from 4,624 reviewed SwissProt proteins (Guo et al. 2015). The library was compiled using DDA data of the kidney tissues in the same mass spectrometer. Protein isoforms and protein groups were excluded from this analysis. The peptides identified were aligned prior to protein inference using the algorithm TRansition of Identification Confidence (TRIC) (version r238), which is available from https://pypi.python.org/pypi/msproteomicstools and https://code.google.com/p/msproteomicstools. The parameters used for the feature_alignment.py program are: max_rt_diff=30, method=global_best_overall, nr_high_conf_exp=2, target_fdr=0.001, use_score_filter=1. The two most intense peptides were used to quantify proteins. 3,123 proteins were quantified across all samples with peptide and protein FDR below 1%.

### Protein complexes (Subnets)

Subnets can be determined *a priori* and independent of the data used for analysis. For example, decomposition of a network into subnets can be optimized via functional coherence evaluation (Goh et al. 2012c). However, complexes are true biological subnets, and proven to be superior to inferred ones (Goh et al. 2013b). They are also stable as they are determined independently of the proteomic data. Thus the complex-based feature vector can be used in generalizability studies comparing related genomic and proteomic data.

Protein complexes were obtained from CORUM database which contains manually annotated protein complexes from mammalian organisms (Ruepp et al. 2008). These are considered gold-standard subnets based on our previous work (Goh et al. 2013b). Although the original algorithms recommended subnets of at least size 5, it is difficult for proteomics to meet this criteria. As a compromise, to keep in line with the original requirements of RBNA algorithms (Soh et al. 2011; Lim and Wong 2014), we considered protein complexes with at least 3 components that were identified and measured in the proteomics screen.

### Standard protein-based feature selection using t-test (SP)

As a control to why network methods are required to extend proteomic analysis, a t-statistic (*T*_*p*_) is calculated for each protein *p* by comparing the z-normalized expression scores between classes *C1* and *C2*, with the assumption of unequal variance between the two classes (Raju 2005).

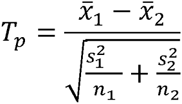

where 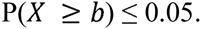 is the mean expression level of the protein *p*, *s*_*j*_ is the standard deviation and *n*_*j*_ is the sample size, in class *Cj*.

The *T*_*p*_ is compared against the nominal t-distribution to calculate the corresponding p-value. A feature is deemed significant if p-value ≤ 0.05.

### Hypergeometric Enrichment (HE)

HE is a standard hypergeometric enrichment pipeline performed in many earlier studies (Goh et al. 2012a) and consists of 2 parts: first, differential proteins are identified using the unpaired two-sided t-test between normal and disease samples using their z-normalized protein expressions (This is similar to SP) (Rivals et al. 2007). Proteins with p-value ≤ 0.05 are considered differential.

In the second part, this is followed by a hypergeometric enrichment analysis against the protein complexes (p-value ≤ 0.05). Given a total number of proteins *N*, with *B* of these belonging to a complex and *n* of these proteins in the test set, the probability *P* that *b* or more proteins from the test set are associated by chance with the complex is given by:

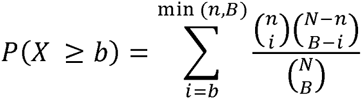

The complex is deemed significant in HE if P(*X* ≥ *b*) ≤ 0.05.

### SNET/FSNET/PFSNET/PPFSNET

The RBNAs (SNET, FSNET, PFSNET and PPFSNET) are similar algorithms but differing in certain key assumptions or test set-ups. The first three RBNAs can be seen as improvements on top of each other. PPFSNET, which we are proposing as a “paired-sample” version of PFSNET, is introduced here for the first time.

We begin with a description of SNET:

Given a protein *gi* and a tissue *pk*, let *fs*(*gi*,*pk*) = 1, if the protein *gi* is among the top alpha percent (default = 10%) most-abundant proteins in the tissue *pk*; and = 0 otherwise.

Given a protein *gi* and a class of tissues *Cj*, let

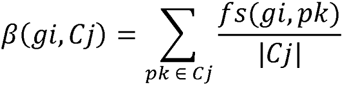

That is, *β*(*gi*,*Cj*) is the proportion of tissues in *Cj* that have *gi* among their top alpha percent most-abundant proteins.

Let *score*(*S*,*pk*,*Cj*) be the score of a protein complex *S* and a tissue *pk* weighted based on the class *Cj*. It is defined as:

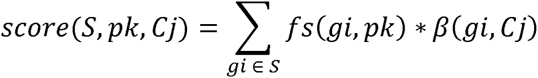

The function *f*_*SNET*_(*S*,*X*,*Y*,*Cj*) for some complex *S* is a t-statistic defined as:

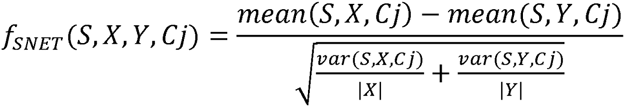

where *mean*(*S*,#,*Cj*) and *var* (*S*,#,*Cj*) are respectively the mean and variance of the list of scores {*score*(*S*,*pk*,*Cj*) | *pk* is a tissue in # }.

The complex *S* is considered significantly highly abundant (weighted based on *Cj*) in *X* but not in *Y* if *f*_*SNET*_(*S*,*X*,*Y*,*Cj*) is at the largest 5% extreme of the Student t-distribution, with degrees of freedom as determined by the Welch-Satterwaite equation.

Given two classes *C1* and *C2*, the set of significant complexes returned by SNET is the union of {*S* | *f*_*SNET*_(*S*,*C1*,*C2*,*C1*) is significant} and {*S* | *f*_*SNET*_(*S*,*C2*,*C1*,*C2*) is significant}, the former being complexes that are significantly consistently highly abundant in *C1* but not *C2*, the latter being complexes that are significantly consistently highly abundant in *C2* but not *C1*.

FSNET is identical to SNET, except in one regard:

For FSNET, the definition of the function *fs*(*gi*,*pk*) is replaced so that *fs*(*gi*,*pk*) is assigned a value between 1 and 0 as follows: *fs*(*gi*,*pk*) is assigned the value 1 if *gi* is among the top alpha1 percent (default = 10%) of the most-abundant proteins in *pk*. It is assigned the value 0 if *gi* is not among the top alpha2 percent (default = 20%) most-abundant proteins in *pk*. The range between alpha1 percent and alpha2 percent is chopped into *n* equal-sized bins (default =4), and *fs*(*gi*,*pk*) is assigned the value 0.8, 0.6, 0.4, or 0.2 depending on which bin *gi* falls into in *pk*.

A test statistic *f*_*FSNET*_ is then defined analogously to *f*_*SNET*_. Given two classes *C1* and *C2*, the set of significant complexes returned by FSNET is the union of {*S* | *f*_*FSNET*_(*S*,*C1*,*C2*,*C1*) is significant} and {*S* | *f*_*FSNET*_(*S*,*C2*,*C1*,*C2*) is significant}.

For PFSNet, the same *fs*(*gi*,*pk*) function as in FSNet is used. But it defines a score *delta*(*S*,*pk*,*X*,*Y*) for a complex *S* and tissue *pk* wrt classes *X* and *Y* as the difference of the score of *S* and tissue *pk* weighted based on *X* from the score of *S* and tissue *pk* weighted based on *Y*. More precisely: *delta*(*S*,*pk*,*X*,*Y*) = *score*(*S*,*pk*,*X*) – *score*(*S*,*pk*,*Y*).

If a complex *S* is irrelevant to the difference between classes *X* and *Y*, the value of *delta*(*S*,*pk*,*X*,*Y*) is expected to be around 0. So PFSNet defines the following one-sample t-statistic:

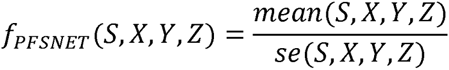

where *mean*(*S*, *X*, *Y*, *Z*) and *se*(*S*, *X*, *Y*, *Z*) are respectively the mean and standard error of the list {*delta*(*S*,*pk*,*X*,*Y*) | *pk* is a tissue in *Z*}. The complex *S* is considered significantly consistently highly abundant in *X* but not in *Y* if *f*_*PFSNet*_(*S*, *X*, *Y*, *X* ∪ *Y*) is at the largest 5% extreme of the Student t-distribution.

Given two classes *C1* and *C2*, the set of significant complexes returned by PFSNet is the union of {*S* | *f*_*PFSNet*_(*S*,*C1*,*C2*,*C1* ∪ *C2*) is significant} and {*S* | *f*_*PFSNet*_(*S*,*C2*,*C1*,*C1* ∪ *C2*) is significant}, the former being complexes that are significantly consistently highly abundant in *C1* but not *C2*, the latter being complexes that are significantly consistently highly abundant in *C2* but not *C1*.

The above formulation of PFSNet is for the situation where tissues in *C1* and *C2* are unpaired. If paired tissues are used, a paired-sample version of PFSNet (PPFSNET) can be formulated as follows.

Given a subject *pk*, we write *pkA* to denote his tissue in class *C1* and *pkB* to denote his paired tissue in class *C2*. Then we define the following paired delta score of the complex *S* and subject *pk* wrt classes *X* and *Y*:

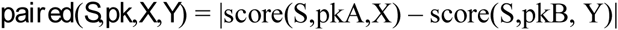

If the complex *S* is irrelevant to the difference between classes *X* and *Y*, as mentioned earlier, then the mean of *paired*(*S*,*pk*,*X*,*Y*) is expected to be 0. We define a one-sample t-statistic to test for this:

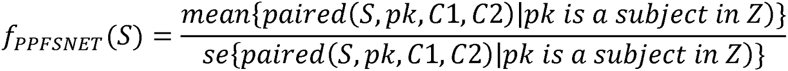

where *Z* is the set of all subjects with paired tissues in *C1* and *C2*. A complex S is considered significant, if f__PPFSNET_(S) is at the largest 5% extreme of the Student t-distribution.

### Performance Benchmarks

We propose a set of 3 criteria for the performance benchmarks. These are precision/recall, feature-selection stability and cross-validation predictive accuracy.

**Precision/Recall** --- In precision/recall, the significant complexes *c*, from each subsampling simulation is benchmarked against the total set of significant complexes, *C*, derived from an analysis of the complete dataset. We make the assumption that the complete dataset is representative of the population. Thus, a completely precise method based on a subsampling should report a subset *c* of *C* (*c* ⊆ *C*) as significant, and no more (considered false positives). Similarly, perfect recall should report all complexes in *C* (i.e., *c* = *C*) as significant.

We can calculate precision and recall as follows:

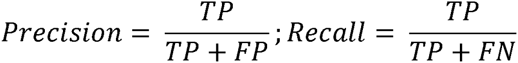

where TP, FP and FN are the True Positives, False Positives and False Negatives respectively.

Both measurements are important in evaluating the performance of a method. E.g., a method that is precise but not sensitive would make some good-quality predictions but may not provide enough data for model building or understanding the phenomena. A highly imprecise but sensitive method may capture all relevant features but at the cost of introducing much noise (irrelevant features). A good method must be both precise and sensitive. There are several metrics for combining precision and sensitivity; an example is the F-score (*F*_*S*_):

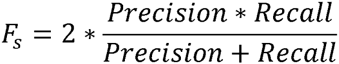

These evaluation metrics require prior knowledge of the set of TP, TN, FP and FN in the dataset. In biology and particularly in proteomics, such “gold standard” data does not exist. We therefore make the assumption that the full dataset is the population, and apply the given feature-selection algorithm to determine the total set of TPs. Comparisons of features selected by repeated subpopulation sampling against those from the complete dataset will provide us a sense of the precision and recall rates. However, we must highlight a caveat to this way of defining the gold standard: It may mislead when the given feature-selection algorithm is unstable, as the algorithm is likely to return an entirely different “gold standard” set of features when applied to a different dataset.

**Feature-selection stability** --- A good feature-selection method must be able to make consistent and reproducible selections, even at small sample sizes. Across different samplings, the technique should reliably provide similar findings. A method with generally high accuracy but low stability has limited utility. It is well known that depending on the dataset, or different parts of the dataset, the same test can select highly different feature sets; this can be attributed to a lack of statistical power and/or using unreliable p-value.

For each method, we took random samplings of size 4, 6 and 8 tissues from both normal and cancer classes (n=12) to simulate small (4) to moderate (8) sample size scenarios. This is repeated 1,000 times to generate a binary matrix, where each row is a simulation, a value of 1 indicates a complex is significant, and 0 otherwise.

The binary matrix is used for comparing stability and consistency of significant features produced by each method. Three evaluations on the binary matrix can be performed: 1/ row-wise comparisons based on the Jaccard coefficient to evaluate cross-simulation similarity, 2/ row summation to evaluate the distribution of the number of selected significant complexes, and 3/ column summation to evaluate the persistence and stability of each selected significant complex.

The feature-***selection stability score is calculated as follows:*** The columns in the binary matrix generated per bootstrap sampling represent the all complexes being tested, while the rows represent the number of simulations (n=1000). A value of 1 means the complex turned up significant while 0 means it did not. Summing each column and dividing it by the number of simulations provides a single stability vector containing the normalized values indicative of complex stability (0 means the complex was never observed, while 1 means the complex was significant across all 1000 simulations).

To calculate a unified score for feature-selection stability, first, all 0 values are discarded from the stability vector (since these are complexes that have never been observed even once across all simulations, and thus irrelevant). Next, the remaining values are summed and divided by the total length of the stability vector, thus generating the feature-selection stability score.

**Cross-validation prediction accuracy** --- Predictive accuracy in the cross-batch scenario considers whether the selected features have any generalizability to other similar datasets. I.e., if we identify x features in one dataset that can discriminate between classes *C1* and *C2*, can these x features do well in another? Unfortunately, we do not have a second batch of data. Thus, we performed a cross-validation study instead. Specifically, we split the data equally into 2 sets (training and testing; n=12 each) and performed feature selection using each method. We trained a Naïve Bayes classifier on the significant features identified from training set, and measured accuracy of the trained classifier on the test set, where

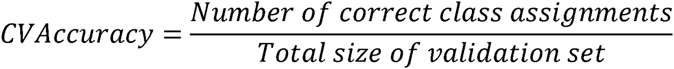

This was repeated 10 times on 10 different random splits of the data into training and testing sets.

To determine if the CV accuracy is meaningful, we randomly picked an equal number of features 1,000 times and retrained the Naïve Bayes classifier to produce a vector of null accuracy values. The CVAccuracy p-value is the number of times null accuracy ≧ 0.99 over 1,000.

The CV accuracy and the CV p-value can be combined by taking the ratio CV accuracy/ CV p-value. Therefore, a method with high accuracy and low p-value will generate a higher score.

## Results and discussions

### Single Protein (SP)-based and hypergeometric-based enrichment (HE) are flawed approaches and unsuitable for clinical applications

SP has better feature reproducibility and feature-selection stability than HE. Based on the distributions of the Jaccard coefficients, SP has higher inter-sampling agreement rates than HE (Figure 1A) --- i.e., the median Jaccard coefficients are higher, with smaller inter-quantile range. While SP benefits from increased sampling size, it is noteworthy that HE’s inter-sampling similarity rates does not improve likewise. This suggests that HE is unstable.

**Figure 1.**
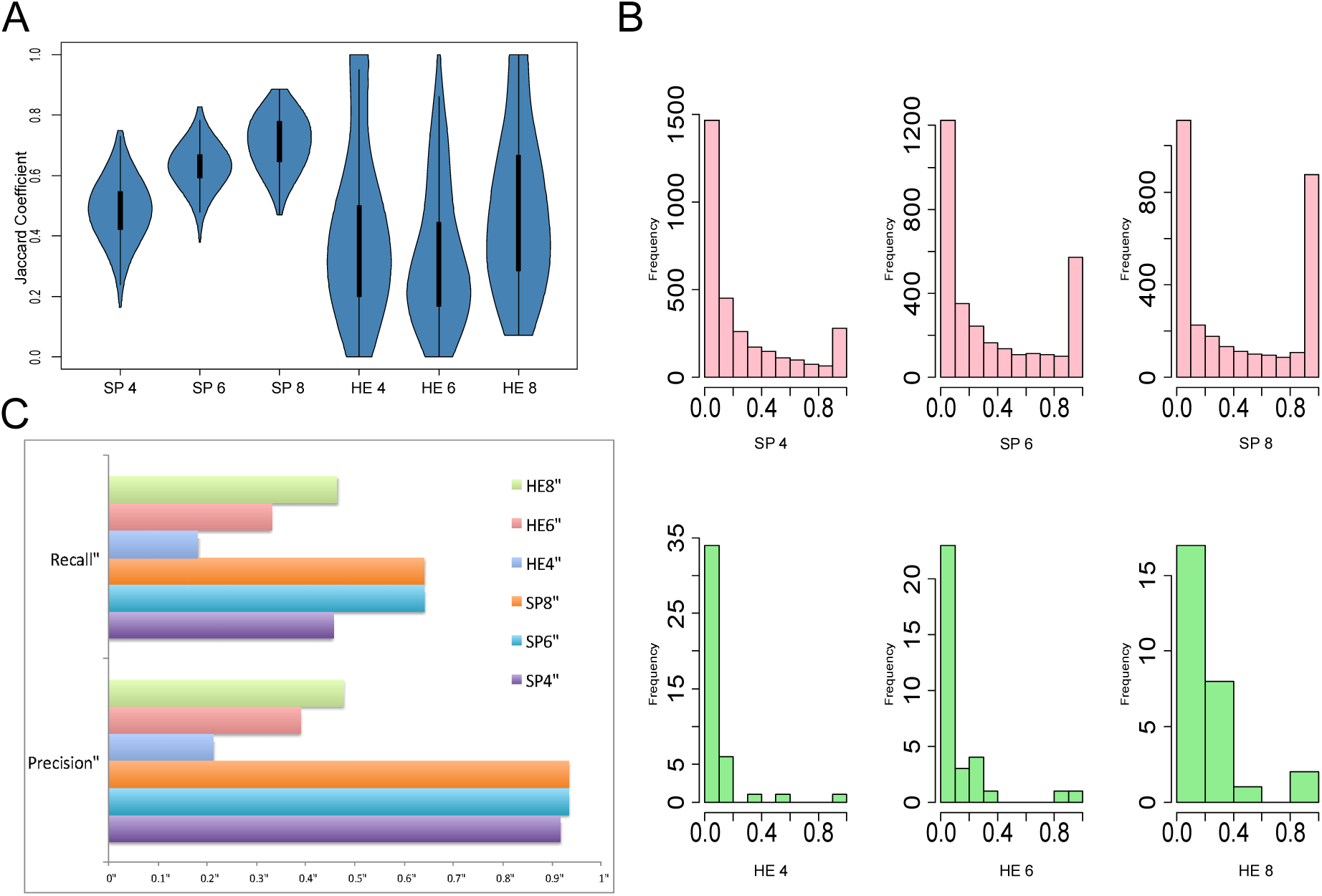
Single Protein (SP) and Hypergeometric Enrichment (HE) feature-selection stability and precision-recall rates. **A: Pairwise feature vector similarity across random samplings.** SP and HE were evaluated 1,000 times on random subsets of sizes 4, 6 and 8. Simulations were compared pairwise for reproducible features using the Jaccard Coefficient. SP benefits from increased sampling size and is relatively more stable than HE. **B: Feature-selection stability across 1,000 simulations**. With increased sampling size, both SP and HE’s feature-selection stability improved, i.e., an observable right shift in the histograms. In both cases however, a vast majority of selected features was never consistently reproduced. **C: Precision/Recall Plots**. Across 1,000 resamplings at sizes 4, 6 and 8, the significant features were compared against the features identified using the full dataset. SP has higher precision than HE. But both HE and SP suffer from relatively modest recall rates.

Likewise, as sampling size increases, the proportion of stable proteins increases in SP (Figure 1B; pink). It is particularly interesting while SP’s feature-selection stability does improve with sampling size, as reflected by the increased number of features with stability scores of 1. A large proportion of proteins still remain unstable (close to 0). This suggests that many of the SP-significant proteins have low stability, and are therefore poorly reproducible. In HE, increasing sampling size has lower effects on overall feature-selection stability score improvements. Note that in HE, the features are complexes.

Based on the same sampling simulations (n= 4 to 8), SP exceeds HE both in precision and recall rates (Figure 1C). This can be attributed to HE’s low feature-selection stability. On the other hand, SP has very high precision rates while maintaining reasonable recall.

In SP, when the t-test deployed is applied on full data, more than half the proteins turned up significant (1649 out of 3123; ∼the DOF is essentially on5e3%) at the 95% significance level. Intuitively, we would expect that many of these are false positives.

To identify the potential cause of the high false-positive rate (FPR), we randomly assigned the normal samples into two groups 1,000 times, and performed the t-test on each protein. The resultant FPR appears reasonable, with a median of 44 (E-value = 1000 * 0.05 = 50), and does not account for the high number of hits in the normal-vs-cancer situation.

We repeated a similar analysis by randomizing the cancer samples, and found that the FPR is only slightly higher with a median of 65. Clearly, the high FPR in the cancer-vs-normal scenario cannot be attributed to extreme heterogeneity within each class.

Comparisons of the distribution of log expressional scores did not suggest that one class has consistently higher expression than the other (Supplementary Figure 1A). The respective averaged means for normal and cancer classes are -0.03 and 0.03 (Supplementary Figure 1B). Thus, the expression center for cancer samples is slightly higher than normal samples. Despite this small difference, in our observation, this may be correlated to the high FPR, which supports the notion that the standard t-test for proteins based on expression data is indiscriminative.

Since more than half of the proteins are significant, it is unsurprising that SP precision rates are high. The random samplings generate subsets of the SP significant proteins, and thus, lower the recall rates accordingly.

The reproducibility and reliability of the p-value is a critical concern often overlooked in analysis (Halsey et al. 2015). Since the t-test returns a large number of proteins (1649) at the 95% significance level, we expect the specificity of the test to be low, and therefore high sample-to-sample variability for the p-value computed for the same feature. We randomly sampled 5 to 200 out of 1,649 proteins originally reported as significant in the full dataset. We recomputed the t-test p-values in half of the dataset, and computed the mean of the p-values. This was performed 1,000 times per sampling size (Figure 2A). Although the variability of the p-value means expectedly decreases as sampling size increases, the median hovers high above the 5% significance level, suggesting that many of the 1,649 significant proteins are in fact, unreliable features.

**Figure 2.**
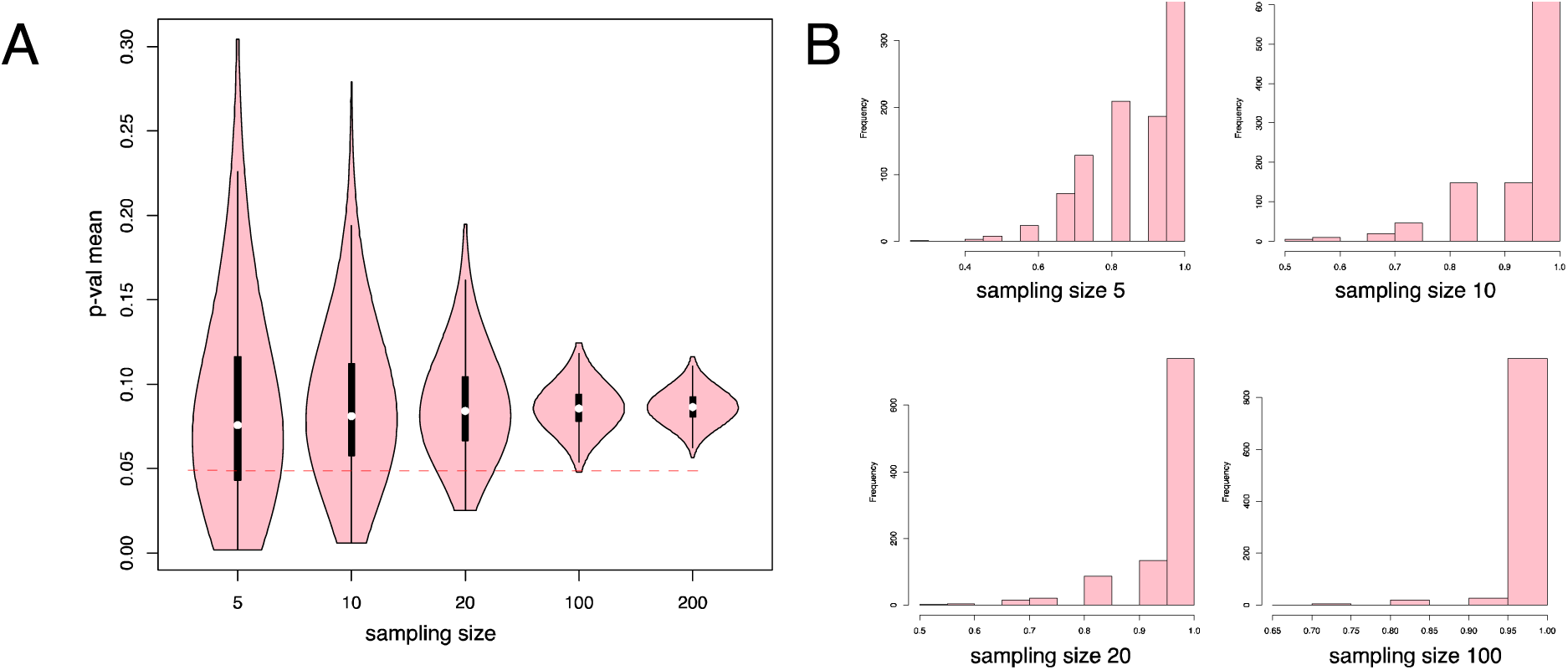
SP has low reproducibility and cross-validation A: T-test p-value means distribution recalculated on half the original dataset. 1,649 proteins were significant on the full dataset in SP. Random subsets of these were chosen, and the p-value means recalculated on half the original data. When n is small, the variability of the p-value means is expectedly high. As n increases, the variability of p-val means decreases, but the median line falls above the 0.05 line. This suggests that a good number of features have unstable p-values, and their significance values cannot be reliably reproduced. **B: SP’s Cross Validation (CV) accuracy is statistically insignificant.** Only a small number of SP-significant features are needed to generate highly accurate Naïve Bayes models. This set of histograms suggests offhand that the high CV accuracy observed in SP (and therefore, HE) has no better predictive accuracy than a cardinal set of randomly picked proteins (Table 1B).

SP has very high cross-validation (CV) accuracy (Table 1). Given that many of the predicted features are unstable and likely false positives, this finding also warrants closer inspection. We found that any random combination of features (even small combinations at n=5) would give very high CV accuracies anyway (Figure 2B). To provide statistical confidence for the observed CV accuracy, we computed a CV p-value based on the null distribution generated by sampling equal sized protein sets 1,000 times. We found that the CV p-values for SP were consistently above 0.9, and thus insignificant (Table 1B). HE uses the same null distribution as SP (since there is no formal way of scoring complexes in HE, the SP-significant proteins in the HE-significant complexes are used as HE features in this cross-validation study). Since HE picks on average 10x less proteins than SP, theoretically, it is expected it should have better CV p-values. However, the null distributions saturate close to n=100 (most of the null accuracies are 1) (Figure 2B). Thus, the CV p-values in HE are similar to those for SP (Table 1B).

**Table 1.** A: Cross-Validation (CV) accuracy of Naïve Bayes models in Single Proteins (SP) and Hypergeometric Enrichment (HE). HE is a subset of SP (about ten times less), but both share seemingly high prediction accuracy. **B: Estimated CV p-values for SP and HE**. Despite high CV accuracy, the SP and HE results perform no better than randomly selected gene sets. The distribution of the null model is saturated around 100 randomly picked genes (cf. Figure 2A). Thus, even though SP is usually larger by a factor of 10 than HE, the CV p-values are similar.

| Group | SP |  |  | HE |  |  |
| --- | --- | --- | --- | --- | --- | --- |
|  | No. significant features (0.05) | self-validation | cross-validation | No. significant features (0.05) | self-validation | cross-validation |
| 1 | 901 | 1.00 | 1.00 | 92 | 1.00 | 1.00 |
| 2 | 1213 | 1.00 | 1.00 | 174 | 1.00 | 1.00 |
| 3 | 908 | 1.00 | 1.00 | 80 | 1.00 | 1.00 |
| 4 | 1210 | 1.00 | 1.00 | 213 | 1.00 | 1.00 |
| 5 | 800 | 1.00 | 1.00 | 126 | 1.00 | 1.00 |
| 6 | 1825 | 1.00 | 0.75 | 358 | 1.00 | 0.75 |
| 7 | 1017 | 1.00 | 1.00 | 124 | 1.00 | 1.00 |
| 8 | 1284 | 1.00 | 1.00 | 195 | 1.00 | 1.00 |
| 9 | 1251 | 1.00 | 1.00 | 164 | 1.00 | 1.00 |
| 10 | 834 | 1.00 | 1.00 | 91 | 1.00 | 1.00 |
| mean | 1124.30 | 1.00 | 0.98 | 161.70 | 1.00 | 0.98 |
| s.d. | 306.48 | 0.00 | 0.08 | 82.77 | 0.00 | 0.08 |
| COV | 0.27 | 0.00 | 0.08 | 0.51 | 0.00 | 0.08 |

| SP |  | HE |  |
| --- | --- | --- | --- |
| No. significant features (0.05) | CV pval | No. significant features (0.05) | CV pval |
| 901 | 0.92 | 92 | 0.92 |
| 1213 | 0.91 | 174 | 0.91 |
| 908 | 0.92 | 80 | 0.92 |
| 1210 | 0.91 | 213 | 0.91 |
| 800 | 0.92 | 126 | 0.92 |
| 1825 | 0.89 | 358 | 0.89 |
| 1017 | 0.90 | 124 | 0.90 |
| 1284 | 0.93 | 195 | 0.93 |
| 1251 | 0.92 | 164 | 0.92 |
| 834 | 0.92 | 91 | 0.92 |
| 1124.30 | 0.91 | 161.70 | 0.91 |
| 306.48 | 0.01 | 82.77 | 0.01 |
| 0.27 | 0.01 | 0.51 | 0.01 |

We conclude that despite the seemingly high CV accuracy for SP and HE, it is no better than picking any random set of proteins. This finding is consistent with Venet et al’s observation that, in breast cancer, any set of 100 genes or more selected at random has a 90% chance to be significantly associated with outcome (Venet et al. 2011).

Aside from having similar CV performance to SP, HE has mediocre performance in the other reliability benchmarks. Thus, despite its being commonly used for identifying enriched subnetwork modules, it appears that HE should be avoided.

HE tests complexes based on significant features determined by the same procedure in SP. Hence, it is important to first understand how feature-selection stability (reproducibility), t-test p-value (significance testing) and Bayes weights (cross-validation) are related in SP, and how HE might affect these relationships. Supplementary Figure 2 reveals a positive relationship between stability, t-test p-values and Bayes weights. The proteins with high feature-selection stability have low p-values, and higher assigned Bayes weights. It turns out that HE has a propensity for selecting more stable SP-significant proteins (feature-selection stability determined at sampling n=8 in SP; Figure 1B pink). In Supplementary Figure 3A/3B, the 302 HE-significant proteins (HE SP intersect) are clearly enriched for stable SP-significant proteins.

**Figure 3.**
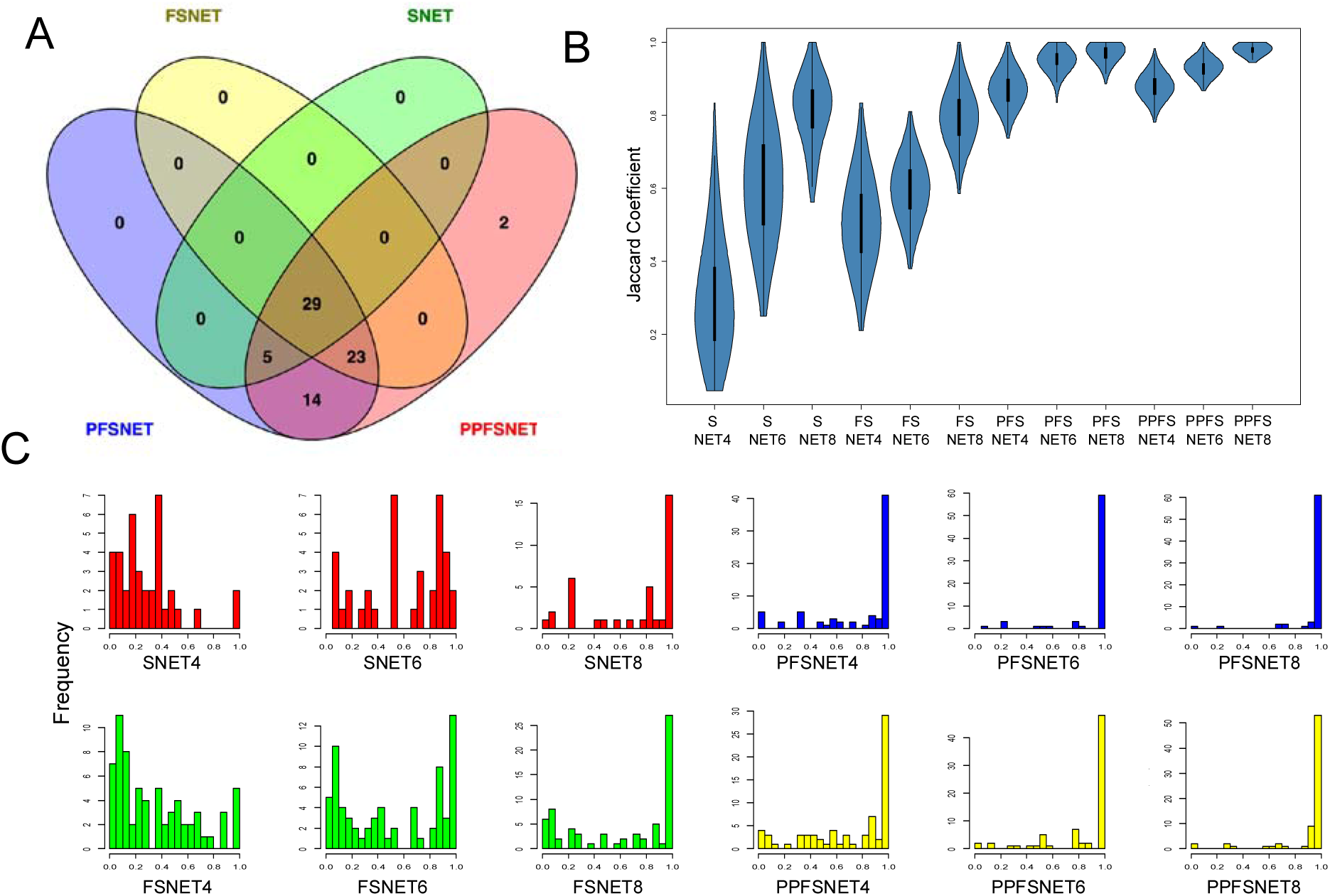
Feature agreements between the 4 ranked-based network approaches (SNet, FSNet, PFSNet and PPFSNET). **A: Significant feature overlaps.** The agreement rates between SNET and FSNet is limited due to the additional incorporation of the alpha2 proteins. PFSNet and PPFSNET are highly sensitive and captured most significant features predicted by the other two. **B: Pairwise global predicted feature similarity across random samplings.** The pairwise feature similarity for SNET, FSNET, PFSNET and PPFSNET are evaluated across random samplings of sizes 4, 6 and 8 (x-axis). The pairwise similarity is calculated as the Jaccard coefficient (intersection/union) and expressed on the y-axis. SNET and FSNET are affected strongly by sample size. PFSNET and PPFSNET on the other hand, are robust against sample size effects. **C: Feature-selection stability distributions.** Unlike SNET (red) and FSNET (green), PFSNET and PPFSNET are very stable even at small sample sizes (x-axis: feature stability scores, y-axis: frequency), with the majority of features consistently reproducible across simulations.

Unfortunately, this beneficial effect of HE is strongly dependent on sample size. We’ve already established that HE has high instability, and therefore the HE hypergeometric p-value has low power. Supplementary Figure 4A confirms this as HE feature-selection stability remains low at the complex and corresponding protein levels even as sampling size increases. There are 302 proteins associated with HE-significant complexes (when tested on the full dataset). Although most of these are enriched for stable SP proteins (Supplementary Figure 3A/3B), Supplementary Figure 4B shows that reasonable enrichment for stable SP proteins can only be achieved at a large sample size (at least 10). Even so, HE cannot capture all stable SP proteins, nor is the SP subset it captures the most statistically significant (Supplementary Figure 3D).

**Figure 4.**
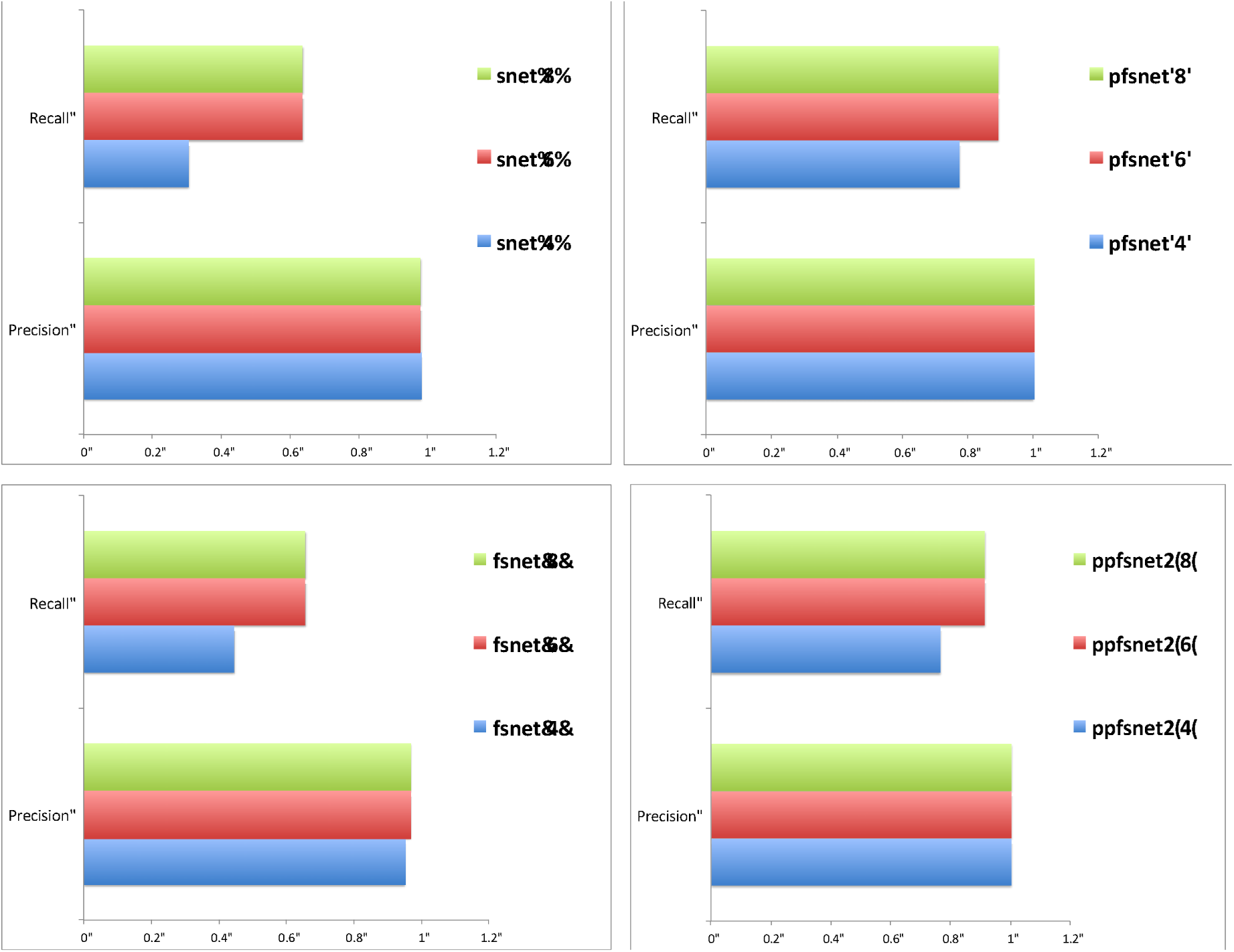
Precision/recall rates between RBNAs (SNET, FSNET, PFSNET and PPFSNET). Across 1,000 resamplings at sizes 4, 6 and 8, the significant features were compared against the full feature set identified using the full dataset. On the whole, the precision rates for all 3 rank-based methods are high. The recall rates for SNET and FSNET are modest, but PFSNET and PPFSNET appear to exhibit very high recall rates while exhibiting similarly high precision rates.

As minimum requirements for identifying biomarker candidates for clinical use, an ideal method should be highly precise and sensitive, stable, and identify relevant features with strong predictive/diagnostic capabilities. Both SP and HE clearly do not meet these requirements.

### Rank-based network methods have high agreement levels and can robustly recover the underlying data classes

Figure 3A shows the overlaps between SNET, FSNET PFSNET and PPFSNET. The results are in agreement with expectation. In earlier transcriptomics analysis, it is known that PFSNET > FSNET > SNET in terms of sensitivity (Lim and Wong 2014).

SNET and FSNET are similar, except with the incorporation of the second alpha level proteins. In line with expectation, most SNET complexes (29/34) were also found among FSNET complexes. FSNET, having considered more proteins, identified an additional 23 complexes.

PFSNET and PPFSNET captured all complexes predicted by SNET and FSNET. Additionally, they agree on an additional 14 complexes not detected by SNET and FSNET. PPFSNET’s paired approach is expected to have higher statistical power than the all-vs-all comparisons in PFSNET, hence its ability to predict more features is expected.

Supplementary Figure 5 shows the Hierarchical Clustering (HCL) heatmaps for predicted features for each method (Euclidean distance, Ward Linkage). Note that there are two sets of matrices per method (matrix weighted by C1, and C2 respectively). In all cases, the cancer and normal classes can be confidently segregated.

**Figure 5.**
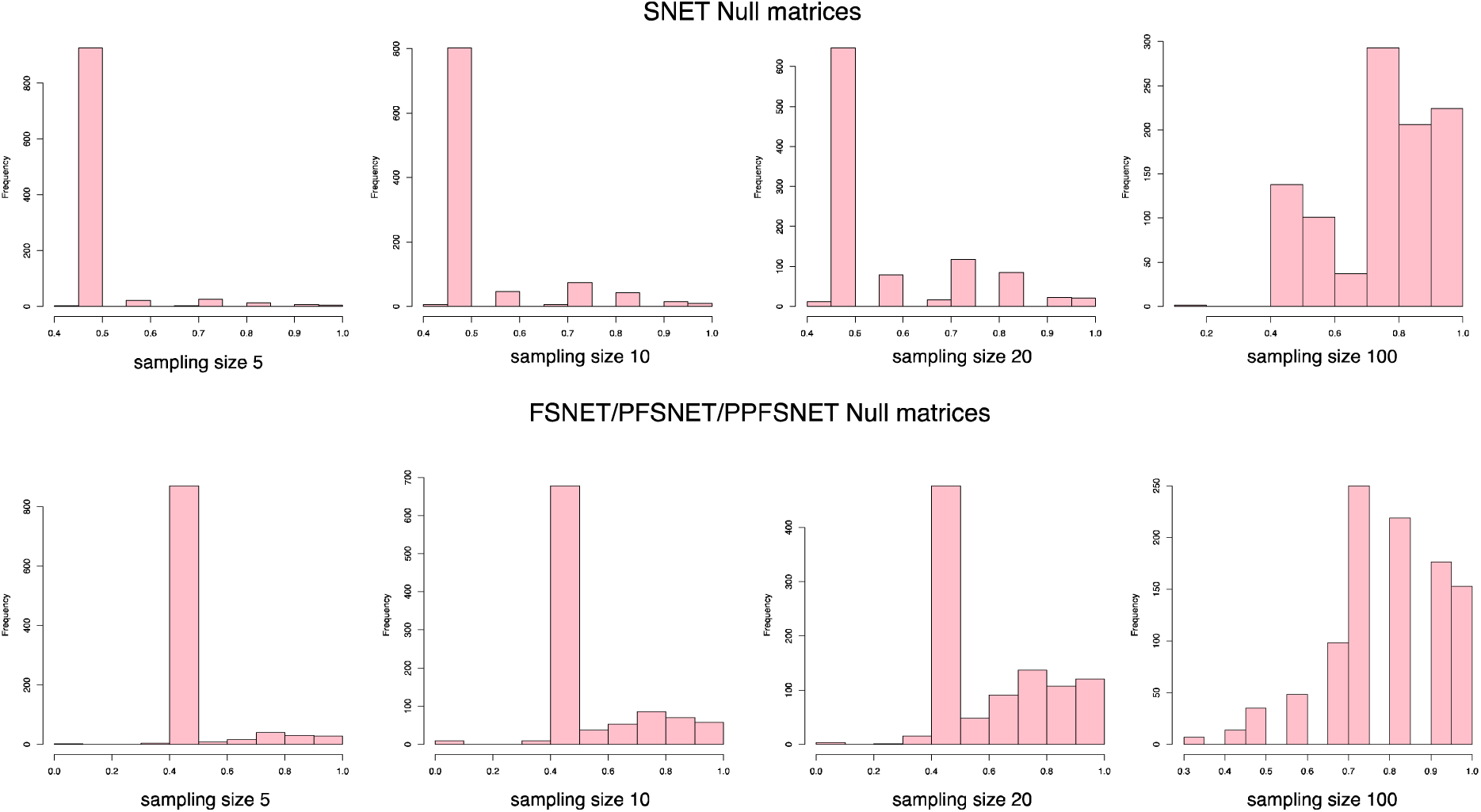
RBNA null models (SNET, FSNET, PFSNET and PPFSNET). The frequency distributions of the CV accuracies (x-axis) for the null are shown here. The null models for the RBNAs are more balanced than SP/HE. SNET and FSNET/PFSNET/PPFSNET null models are plotted separately because the features are scored differently.

### Rank-based network algorithms discover network features with high stability

Figure 3B shows the pairwise prediction similarity between random samplings at sizes 4, 6 and 8 for the RBNAs based on the Jaccard coefficient. PPFSNET and PFSNET dominate in this scenario, with a Jaccard coefficient well above 0.85 even at small sample size. On the other hand, SNET and FSNET are not very stable in the small sample size scenario. While Jaccard coefficient increases appreciably at n=8, at n=4 to 6, the results are comparable to SP (cf. Figure 1A).

For biomarker identification, it is critical that a method provides reproducible biomarkers---i.e., the power of the statistical test is high so that the p-value provided can be relied upon. The Jaccard coefficient provides information on how similar simulation pairs are between samplings globally, but not how stable each selected feature is. To observe this, we calculated the feature stability scores for all complexes over all simulations --- a feature stability score close to 0 means the complex is rarely observed consistently, while a score of 1 means that the feature is always observed. We plotted the frequency distribution of the feature stability scores to observe how stable and reproducible the selections are (Figure 3C).

Again, PFSNET and PPFSNET dominates in feature stability (Figure 3C, yellow and blue) while SNET and FSNET appears less robust, particularly in the small sample size scenario (Figure 3C, red and green. In contrast, Figure 1C shows that SP-and HE-significant features have very low feature stability (pink and light green). In SP, most features were unstable at n=4 (Figure 1C, pink). Although the proportion of stable features increases as n increases, a large proportion of feature stability values are close to 0. HE displays modest improvements as n increases, but the distribution of feature stability values lies on the far left of the histogram, indicating low stability (Figure 1C, light green).

To provide a quantitative means of summarizing the feature stability distributions, the feature-stability scores were averaged over the various sampling sizes, to provide a final score for each method (Table 2A). Here, PPFSNET > PFSNET > SNET > FSNET > SP > HE.

**Table 2.** Summary tables for feature prediction stability and precision/recall. **A: Feature-prediction stability score table.** The feature-prediction stability score vectors are averaged for each sampling size to generate an averaged feature-stability score. Here, PFSNET and PPFSNET are the two best approaches. **B: F-score table.** The F-score is the harmonic mean of precision and recall. PFSNET and PPFSNET have the highest F-scores, indicating they perform well in both precision and recall. It is also of note that PFSNET and PPFSNET can be used in the small sample size scenario, where the F-scores are already high at n=4, and maximized at n=6.

**A**
|  | **Feature-stability scores** |  |  |  |
| --- | --- | --- | --- | --- |
| **Method/Sampling_size** | 4 | 6 | 8 | Avg |
| **SP** | 0.27 | 0.36 | 0.44 | 0.36 |
| **HE** | 0.08 | 0.13 | 0.17 | 0.13 |
| **SNET** | 0.28 | 0.60 | 0.72 | 0.53 |
| **FSNET** | 0.36 | 0.52 | 0.63 | 0.50 |
| **PFSNET** | 0.79 | 0.92 | 0.95 | 0.88 |
| **PPFSNET** | 0.82 | 0.94 | 0.99 | 0.92 |

**B**
|  | **F-scores** |  |  |  |
| --- | --- | --- | --- | --- |
| **Method/Sampling_size** | 4 | 6 | 8 | Avg |
| **SP** | 0.61 | 0.76 | 0.76 | 0.71 |
| **HE** | 0.19 | 0.36 | 0.47 | 0.34 |
| **SNET** | 0.46 | 0.77 | 0.77 | 0.67 |
| **FSNET** | 0.59 | 0.77 | 0.77 | 0.71 |
| **PFSNET** | 0.87 | 0.94 | 0.94 | 0.92 |
| **PPFSNET** | 0.87 | 0.95 | 0.95 | 0.93 |

### Rank-based network algorithms dominate in precision/recall rates

To estimate the RBNA precision/recall rates, we compare the bootstrap resampling simulations against the set of features predicted in the full dataset (population). Figure 4 shows that all RBNAs have high precision rates (close to 1), i.e., the subsamplings predict features that are found in the population, without making many false positive predictions. In fact, sampling size has negligible effects on precision rates.

However, the recall rates are sensitive to sampling size. This is not unexpected --- when sampling size is small, it is less likely that all the causal features will be captured or there is less support for some weaker features, thus lowering recall. SNET and FSNET have relatively low recall rates --- with a peak recall about 0.6. PFSNET and PPFSNET have better recall rates (peak recall ∼ 0.85), which suggest that the statistical tests deployed therein have more power.

We compare this result to Figure 1B where the precision/recall rates for SP and HE are shown. As already shown earlier, SP appears to have very high precision with moderate recall rates. HE on the other hand, has poor precision and recall rates.

The F-score is the harmonic mean of precision and recall, and provides a convenient means of taking both measurements into account simultaneously (Table 2B). The averaged F-scores across sampling sizes (n=4, 6 and 8) are used to evaluate the overall precision/recall performance. Here, PPFSNET > PFSNET > FSNET ties with SP > SNET > HE.

### Rank-based network algorithms provide good Cross-Validation(CV) prediction accuracy

Like SP and HE, RBNAs also perform very well in CV accuracy (Table 3A). However, we also need to consider whether the CV accuracies are statistically significant. Unlike the SP/HE CV accuracy null distributions, the RBNA CV accuracy null distributions are well balanced (Figure 5). Thus, any small number of randomly sampled features based on the RBNA feature matrices will not yield unusually high CV accuracies. As before, to determine a p-value for the RBNA CV accuracy, we sampled complex sets (equal to the number of observed complexes) 1,000 times to generate the null distribution. Table 3B shows that the corresponding CV p-values are substantially lower than those of SP and HE (Table 1B).

**Table 3.** Cross-Validation (CV) accuracy of Naïve Bayes models in SNET, FSNET, PFSNET and PPFSNET. The models were built using the feature scores of the significant features for each method. All methods exhibit high CV accuracy, despite the smaller number of features predicted (much fewer complexes than individual proteins). FSNet has the highest predictive accuracy but is less sensitive than PFSNET or PPFSNET, which identified more complexes. SNET made the least identifications; its sensitivity may be limited by its consideration of only the top 10% proteins.

**A**
| Group | SNET |  |  | FSNet |  |  |
| --- | --- | --- | --- | --- | --- | --- |
|  | No. significant features (0.05) | self_validation | cross_validation | No. significant features (0.05) | self_validation | cross_validation |
| 1.00 | 21 | 0.92 | 0.92 | 34 | 1.00 | 1.00 |
| 2.00 | 24 | 1.00 | 0.92 | 40 | 1.00 | 1.00 |
| 3.00 | 20 | 0.83 | 0.83 | 34 | 1.00 | 1.00 |
| 4.00 | 17 | 1.00 | 0.50 | 38 | 1.00 | 1.00 |
| 5.00 | 26 | 1.00 | 0.92 | 39 | 1.00 | 1.00 |
| 6.00 | 19 | 0.92 | 0.67 | 41 | 1.00 | 0.75 |
| 7.00 | 16 | 0.92 | 1.00 | 33 | 1.00 | 1.00 |
| 8.00 | 19 | 1.00 | 0.67 | 32 | 1.00 | 1.00 |
| 9.00 | 18 | 1.00 | 1.00 | 29 | 1.00 | 0.83 |
| 10.00 | 25 | 0.92 | 1.00 | 37 | 1.00 | 1.00 |
| mean | 20.50 | 0.95 | 0.84 | 35.70 | 1.00 | 0.96 |
| s.d. | 3.44 | 0.06 | 0.17 | 3.89 | 0.00 | 0.09 |
| COV | 0.17 | 0.06 | 0.21 | 0.11 | 0.00 | 0.09 |

| Group | PFSNet |  |  | PPFSNet |  |  |
| --- | --- | --- | --- | --- | --- | --- |
|  | No. significant features (0.05) | self_validation | cross_validation | No. significant features (0.05) | self_validation | cross_validation |
| 1.00 | 65 | 1.00 | 1.00 | 66 | 1.00 | 1.00 |
| 2.00 | 66 | 1.00 | 0.83 | 65 | 1.00 | 1.00 |
| 3.00 | 62 | 1.00 | 1.00 | 66 | 1.00 | 1.00 |
| 4.00 | 69 | 1.00 | 0.92 | 68 | 1.00 | 0.92 |
| 5.00 | 65 | 1.00 | 0.83 | 67 | 1.00 | 1.00 |
| 6.00 | 68 | 1.00 | 0.83 | 69 | 1.00 | 0.83 |
| 7.00 | 67 | 1.00 | 1.00 | 62 | 1.00 | 1.00 |
| 8.00 | 65 | 1.00 | 0.92 | 66 | 1.00 | 0.92 |
| 9.00 | 60 | 1.00 | 0.83 | 61 | 1.00 | 0.92 |
| 10.00 | 63 | 1.00 | 1.00 | 65 | 1.00 | 1.00 |
| mean | 65.00 | 1.00 | 0.92 | 65.50 | 1.00 | 0.96 |
| s.d. | 2.75 | 0.00 | 0.08 | 2.46 | 0.00 | 0.06 |
| COV | 0.04 | 0.00 | 0.09 | 0.04 | 0.00 | 0.06 |

**B**
| SNET |  | FSNET |  |
| --- | --- | --- | --- |
| No. significant features (0.05) | CV pval | No. significant features (0.05) | CV pval |
| 21 | 0.07 | 34 | 0.05 |
| 24 | 0.07 | 40 | 0.07 |
| 20 | 0.05 | 34 | 0.05 |
| 17 | 0.07 | 38 | 0.07 |
| 26 | 0.07 | 39 | 0.07 |
| 19 | 0.06 | 41 | 0.07 |
| 16 | 0.04 | 33 | 0.04 |
| 19 | 0.05 | 32 | 0.04 |
| 18 | 0.04 | 29 | 0.04 |
| 25 | 0.06 | 37 | 0.06 |
| 20.50 | 0.06 | 35.70 | 0.06 |
| 3.44 | 0.01 | 3.89 | 0.01 |
| 0.17 | 0.20 | 0.11 | 0.23 |

| PFSNET |  | PPFSNET |  |
| --- | --- | --- | --- |
| No. significant features (0.05) | CV pval | No. significant features (0.05) | CV pval |
| 65 | 0.07 | 66 | 0.07 |
| 66 | 0.07 | 66 | 0.07 |
| 62 | 0.04 | 66 | 0.07 |
| 69 | 0.07 | 68 | 0.07 |
| 65 | 0.07 | 67 | 0.07 |
| 68 | 0.06 | 70 | 0.07 |
| 67 | 0.04 | 62 | 0.04 |
| 65 | 0.05 | 67 | 0.07 |
| 60 | 0.04 | 64 | 0.07 |
| 63 | 0.06 | 67 | 0.07 |
| 65.00 | 0.06 | 66.30 | 0.06 |
| 2.75 | 0.01 | 2.16 | 0.01 |
| 0.04 | 0.22 | 0.03 | 0.13 |

The CV accuracy and CV p-value are both important and should be considered together. To derive a combined score, we divided the former by the latter (Table 4A) such that a higher score indicates accuracy is high and p-value is low. Using this metric, PPFSNET > FSNET > PFSNET > SNET > SP ties with HE.

**Table 4.** A Summary of cross-validation (CV) accuracies and p-values. CV accuracies and p-values are tabulated for each method. To consider both the accuracy and p-values jointly, we divided the former by the latter such that a larger Acc/P-value ratio is better (higher accuracy, lower p-value). **B Evaluation of methods based on their performance benchmarking**. The sum of ranks shows that PPFSNET is the best method, followed by PFSNET, FSNET, SNET, SP and finally HE.

**A**
| Method | SP | HE | SNET | FSNET | PFSNET | PPFSNET |
| --- | --- | --- | --- | --- | --- | --- |
| CV Accuracy | 0.98 | 0.98 | 0.84 | 0.96 | 0.92 | 0.96 |
| CV Pval | 0.91 | 0.91 | 0.06 | 0.06 | 0.06 | 0.06 |
| Acc/Pval | 1.08 | 1.08 | 14.00 | 16.00 | 15.33 | 16.00 |

**B**
| Method | SP | HE | SNET | FSNET | PFSNET | PPFSNET |
| --- | --- | --- | --- | --- | --- | --- |
| Precision/Recall | 3 | 6 | 5 | 3 | 2 | 1 |
| Feature Stability | 5 | 6 | 3 | 4 | 2 | 1 |
| CV Accuracy/CV p-val | 5 | 5 | 4 | 1 | 3 | 1 |
| Total Ranks | 13 | 17 | 12 | 8 | 7 | 3 |

### PPFSNET is the most robust method

The final summed ranks are summarized in Table 4B. Based on this set of evaluation benchmarks, PPFSNET is the best method, exceling in all 3 benchmarks (total ranks: 3) followed by PFSNET, FSNET, SNET, SP and finally HE.

The key advantage of PPFSNET over PFSNET lies in the statistical test setup, which is only possible if samples can be match-paired --- i.e., normal and cancer tissue originates from same patient. In the paired scenario, inter-sample variability is theoretically reduced, making it more likely to correctly reject the null hypothesis. Hence it is expected that PPFSNET identifies slightly more complexes than PFSNET, but this small increase in feature set translates to stronger performance in all 3 benchmarks. However, the trade-off is in the degrees of freedom (DOF). In the unpaired test, the DOF is essentially one minus-ed off the number of samples, n-1. In the paired-test, because half the samples are used explicitly to compare against the other half, the DOF is n/2 -1.

Generally, RBNAs have high feature-selection stability, precision/recall rates, with a meaningful CV null model while maintaining high CV accuracy rates. PFSNET and PPFSNET are particularly powerful tests and can work even in the small-sample-size scenario, a highly attractive advantage. These RBNAs easily outperform conventional single-protein (SP) expression and hypergeometric enrichment (HE) calculation techniques. However, there is one poignant limitation: because RBNAs work on the rank system, low-abundance proteins are occluded, even if they are indeed informative. While we acknowledge this is a critical limitation, low-abundance proteins generally have poor quantitation reliability; thus, their rank shifts are less informative. Earlier genomic analysis results have proven this point, which resulted in the development of the alpha cut-offs (Soh et al. 2011; Lim and Wong 2014). While networks can provide useful contextualization or impose expressional constraints for the study of important low-abundance proteins, this is beyond the scope of this paper, which explores the suitability of RBNAs as tools for clinical proteomic analysis. A second critical application, which is poorly explored in clinical proteomics, is the ability to isolate and characterize sample sub-populations in highly heterogeneous clinical data. Given that RBNAs have high sensitivity, and can function well even on small samples, they can be attuned towards this, with the expectation of helping develop highly customized treatment methodologies.

## Conclusions

Ranked-based network approaches (RBNAs) extend the utility of proteomic analysis, and increase the likelihood of identifying stable, consistent and generalizable features, even in the small sample size scenario. This in turn, makes it more likely to identify reliable biomarkers/targets in clinical applications.

## Acknowledgements

WWBG is funded by a Professorship of Bioinformatics, School of Pharmaceutical Science and Technology, Tianjin University, China. LW is funded in part by a Singapore Ministry of Education tier-2 grant, MOE2012-T2-1-061.

## Author Contributions

WWBG and LW designed, implemented the bioinformatics method and pipeline, performed analysis, and wrote the manuscript.

## Supplementary Section

### Figures

**Supplementary Figure 1.**
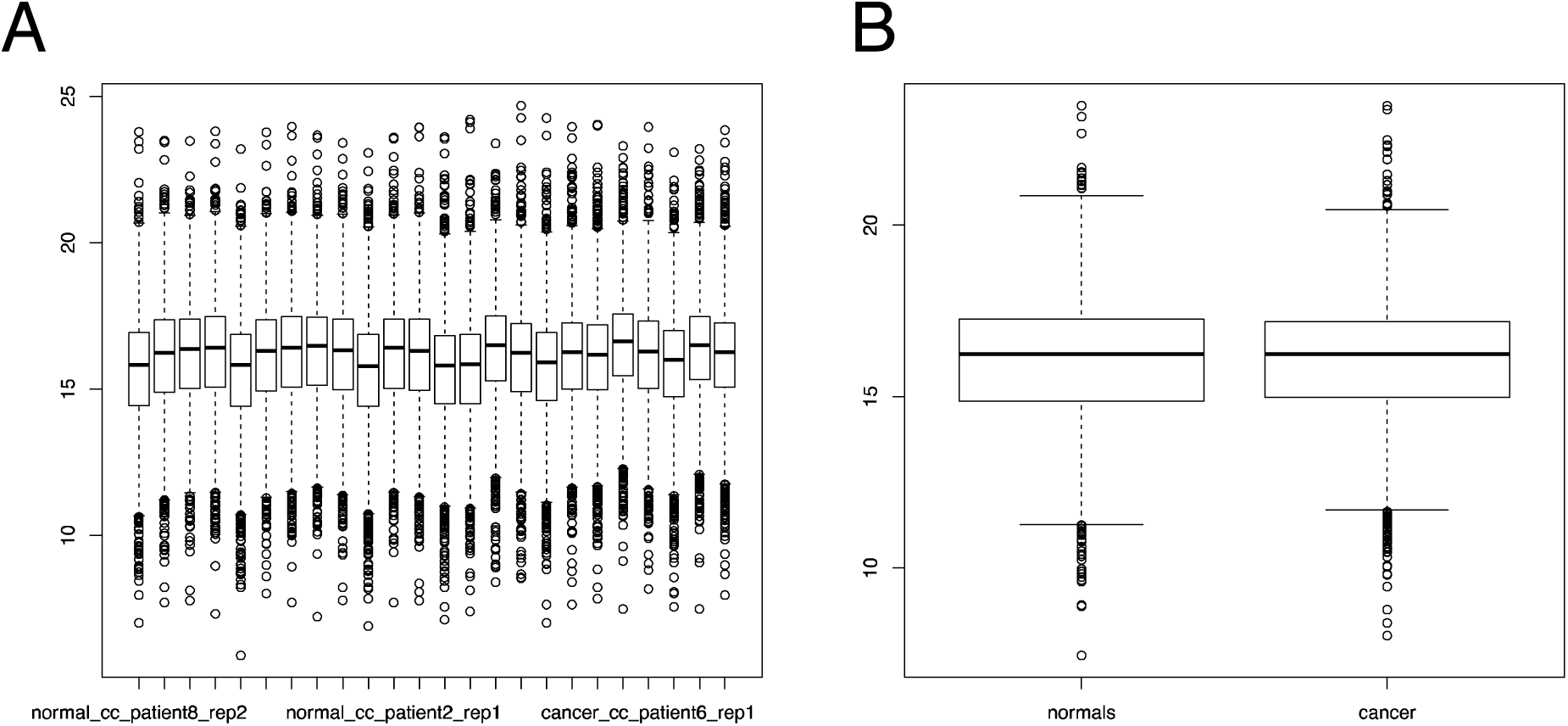
Distribution of log expression scores. A: The distribution of expression values for all samples (across normal and cancer) appears to be even. B: The distribution of expression scores for all proteins averaged by classes normal and cancer. The cancer class has a slightly higher mean than normal. It appears that even subtle differences appear to be able to cause the standard t-test to predict an overly large number of features.

**Supplementary Figure 2.**
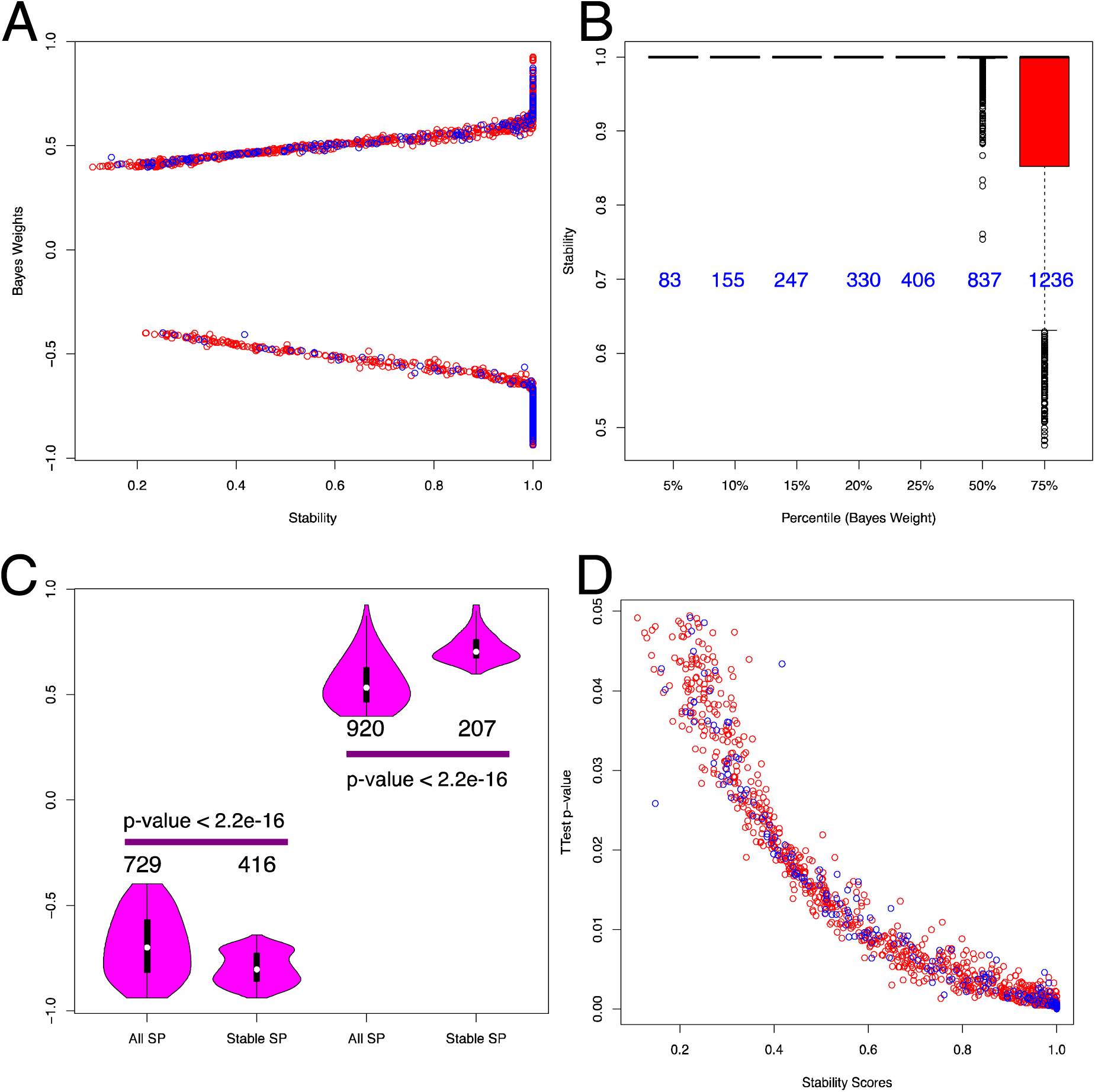
SP proteins that are more stable, have more significant SP p-values and assigned Bayes Weights. A: Bayes weights and stability (feature stability, SP, n=8) are correlated (red --- SP sig, blue --- hyper sig). B: Proteins with high weights are stable. The values in blue indicates the number of proteins at each percentile level. C: Distribution of Bayes weights for all SP sig, and those SP sig with feature stability score of 1 (stable SP). Proteins that are more stable have higher assigned Bayes weights --- Only the Bayes weights with respect to the normal class are used, and consist of both positive and negative weight values. D: Stability scores are correlated to the t-test values. The more stable the score, the more likely it is to have a lower p-value in the SP significant protein set. HE significant proteins are marked in blue.

**Supplementary Figure 3.**
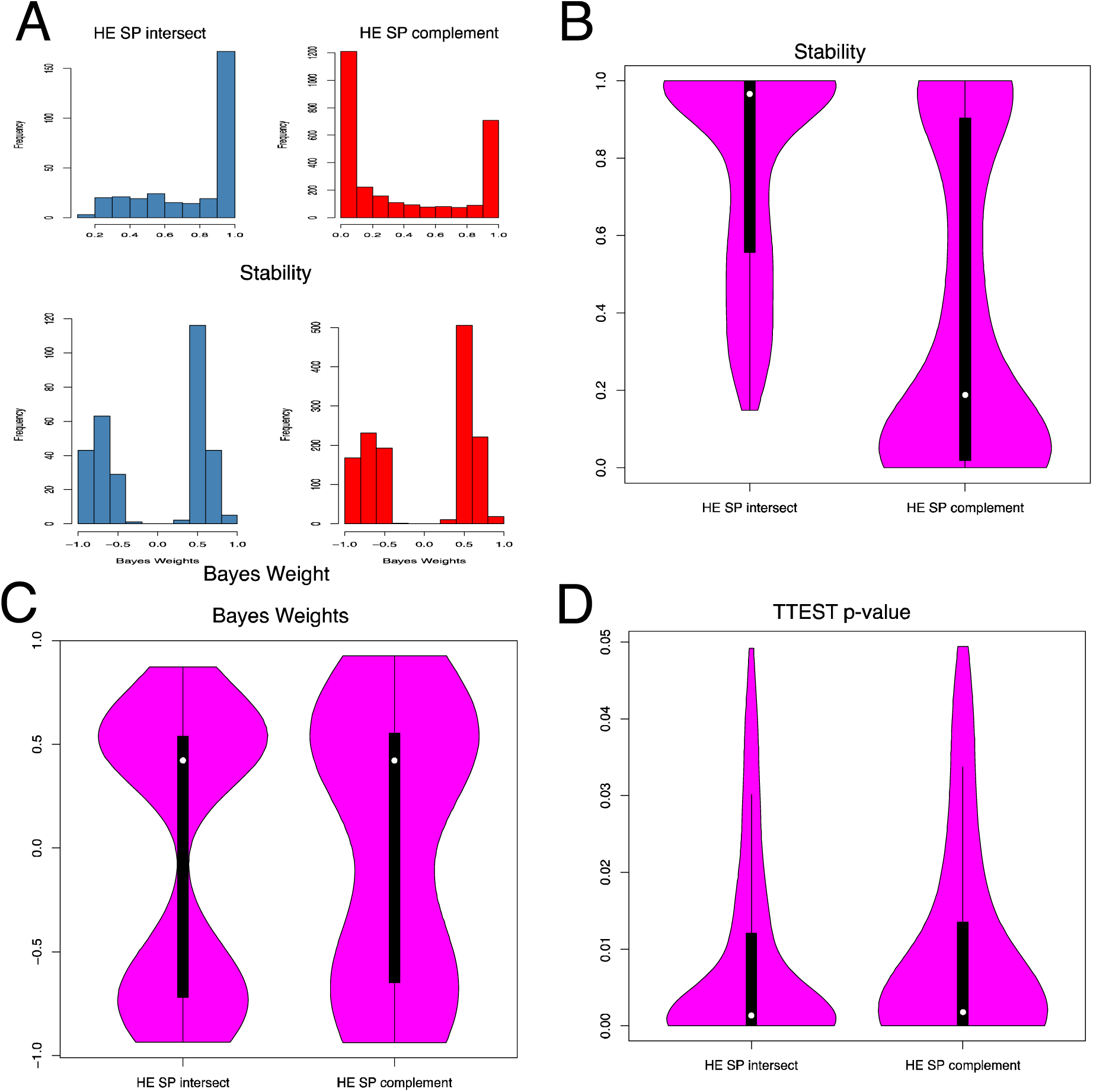
HE selects stable SP proteins. A: Top --- The feature stability distribution of the HE-SP intersection shows that HE tends to select the most stable SP proteins. Bottom --- This propensity to select stable features translates to selecting higher Bayes weights. B and C are elaborations of A showing the violin plots for both Stability and Bayes weights. D: Interestingly, the corresponding p-values are not that different between the intersection and complement. *Note that these observations apply for HE applied on the full dataset.

**Supplementary Figure 4.**
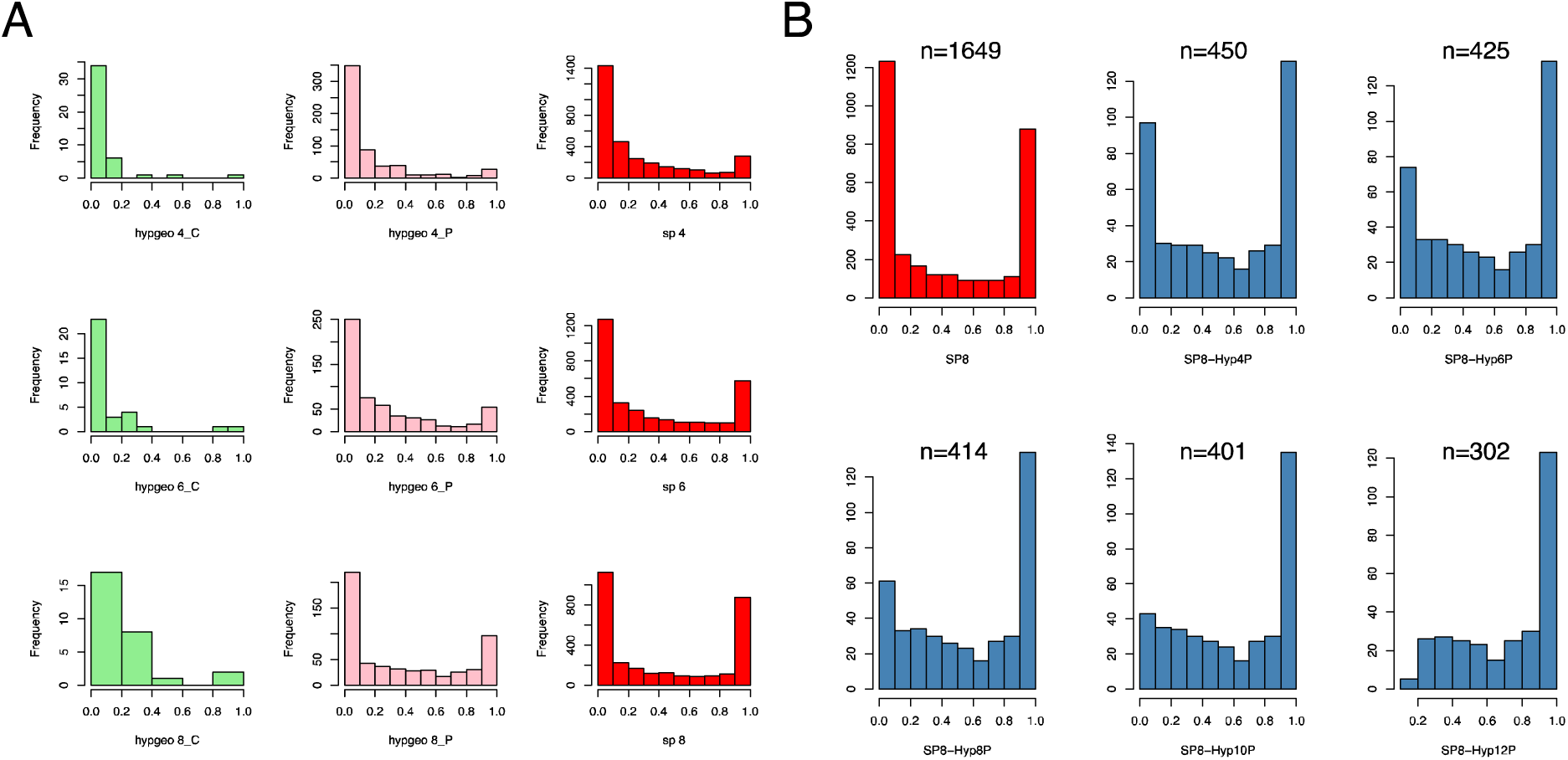
HE’s ability to select stable SP proteins is highly dependent on sampling size. A: hypgeo_n_C and hypgeo_n_P refers to the stability scores for the complexes and the corresponding proteins respectively. Although increasing sampling size doesn’t improve the feature stability at the complex level (green), the feature stability from the corresponding proteins does improve. The SP distribution (red) is shown as a reference. B: Frequency Distribution of feature stability scores at various HE sampling sizes. The histogram in red shows the frequency distribution of feature stability in SP (based on sampling size of 8; SP8). The remaining charts show the intersection between HE and SP significant proteins at various sampling sizes for HE (where Hyp4P, refers to “HE” with sampling size of “4”, and the returned features are the “Proteins” from the significant complexes). The number at the top of each histogram represents the average number of significant HE proteins based on the sampling size. Although HE can potentially select the most stable SP proteins; this ability is dependent on sample size.

**Supplementary Figure 5.**
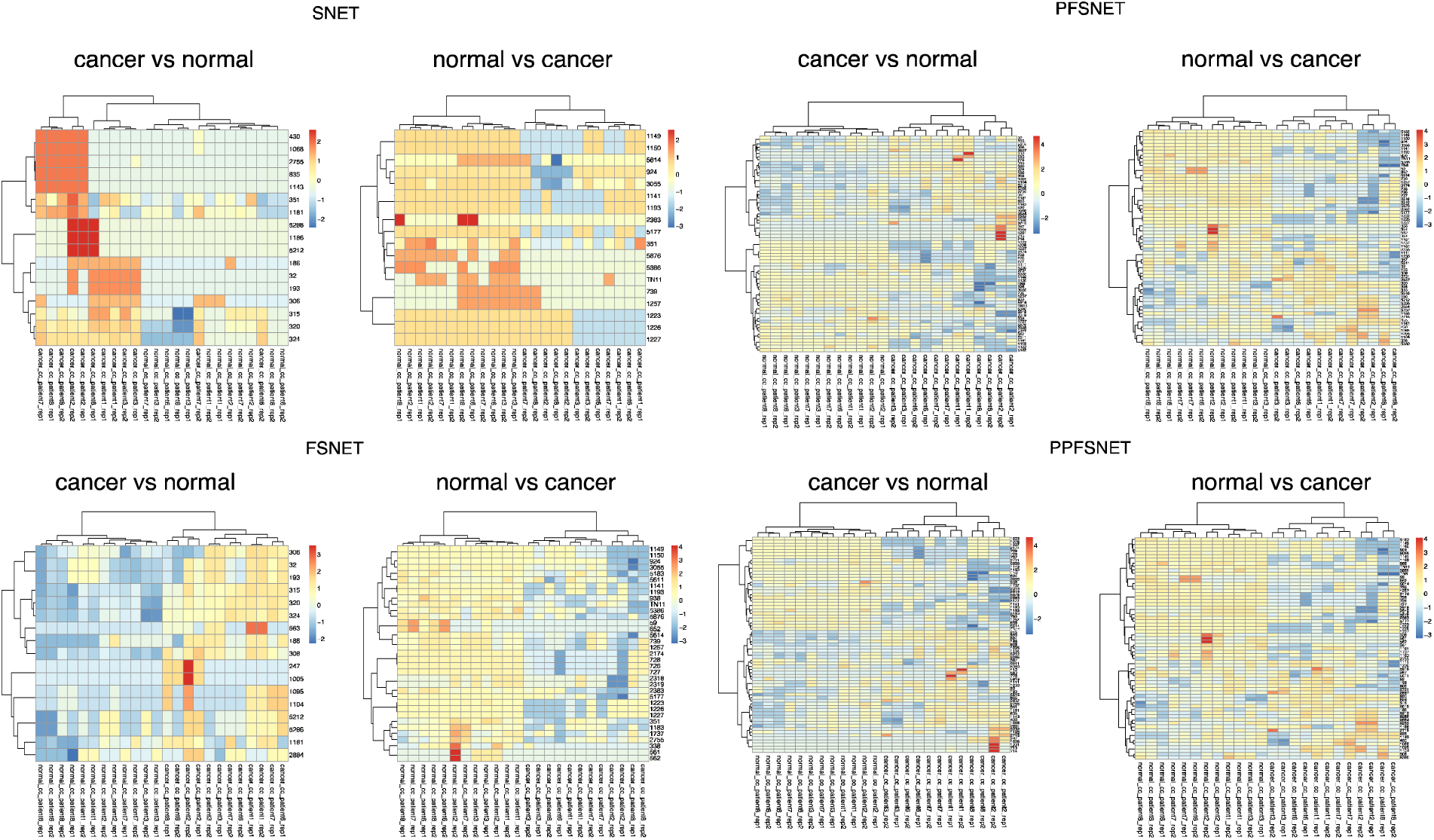
Hierarchical Clustering (HCL) for the RBNAs. The predicted features are highly informative and can recover the underlying classes. A unique attribute of the RBNAs is that two matrices (each weighted by each class) are produced for each dataset. For example, in SNET, for the matrix cancer vs normal, we would be testing for features over-represented for cancer irrespective of the samples classes. And vice versa for normal vs cancer.

